# Aggregation and disaggregation of red blood cells: depletion versus bridging

**DOI:** 10.1101/2024.11.20.624311

**Authors:** Nicolas Moreno, Kirill Korneev, Alexey Semenov, Alper Topuz, Thomas John, Minne Paul Lettinga, Marco Ellero, Christian Wagner, Dmitry A. Fedosov

## Abstract

The aggregation of red blood cells (RBCs) is a complex phenomenon that strongly impacts blood flow and tissue perfusion. Despite extensive research for more than 50 years, physical mechanisms that govern RBC aggregation are still under debate. Two proposed mechanisms are based on bridging and depletion interactions between RBCs due to the presence of macromolecules in blood plasma. The bridging hypothesis assumes the formation of bonds between RBCs through adsorbing macromolecules, while the depletion mechanism results from the exclusion of macromolecules from the inter-cellular space, leading to effective attraction. Existing experimental studies generally cannot differentiate between these two aggregation mechanisms, though several recent investigations suggest concurrent involvement of the both mechanisms. We explore dynamic aggregation and disaggregation of two RBCs using three simulation models: a potential-based model mimicking depletion interactions, a bridging model with immobile bonds, and a new bridging model with mobile bonds which can slide along RBC membranes. Simulation results indicate that dynamic aggregation of RBCs primarily arises from depletion interactions, while disaggregation of RBCs involves both mechanisms. The bridging model with mobile bonds reproduces well the corresponding experimental data, offering insights into the interplay between bridging and depletion interactions and providing a framework for studying similar interactions between other biological cells.

## I. INTRODUCTION

Aggregation of red blood cells (RBCs) is a complex process that strongly affects blood rheology and consequently, plays a crucial role in blood flow and tissue perfusion [1, 2]. Deciphering physical mechanisms that govern RBC aggregation is therefore important for a better understanding of its role in various physiological and pathological processes. RBC aggregation is reversible and influenced by many factors, including the mechanical properties of RBCs, biochemical constituents of blood plasma, and hemodynamic conditions in blood circulation. Among these factors, the role of macromolecules (e.g., fibrinogen, albumin) present in the plasma has been subject to extensive research [3–8].

Two different mechanisms for RBC aggregation have been proposed, including *bridging* and *depletion* [9, 10]. Bridging interaction between RBCs is mediated by macromolecules which adsorb at the RBC surface, and form bridges (or bonds) connecting different cells [11– 13]. Depletion interactions due to the exclusion of macromolecules from the space between RBCs lead to an effective attraction between the cells [14–16]. The depletion model has been partly favored due to numerous studies of RBC aggregation *in vitro*, using solutions with non-blood-borne macromolecules (e.g., dextran) [17–20]. However, both bridging and depletion interactions may occur simultaneously, and depend on the type and concentration of macromolecules. As a result, physical mechanisms that govern RBC aggregation are subject to a long-standing discussion lasting for more than 50 years.

RBC aggregation has been studied by various experimental techniques, including cell aggregation in flow [5, 21– 23], blood sedimentation [24–26], microscopy observations of RBC aggregates [27–30], and the manipulation of cell doublets by optical tweezers [31–34]. Even though most of the experimental methods do not allow us to distinguish between the two different aggregation mechanisms, recent studies with optical tweezers [32, 33] show that forces measured during the dynamic aggregation (i.e., formation) and disaggregation of a RBC doublet can be very different. For pure depletion interactions, no difference in forces responsible for the dynamic aggregation and disaggregation of the doublet should occur, suggesting that different mechanisms for the RBC aggregation are likely at play. Another investigation has explored how the aggregation forces between RBCs are affected, when they are moved between solutions containing different proteins [34]. For purely depletion-mediated interactions, the interaction force between cells is expected to depend only on the concentration of the final solution. However, the experimental results show that the molecular content of the initial solution plays a significant role in determining the RBCs aggregation, providing a further evidence for the existence of bridging interactions. As a result, differences in the forces measured during dynamic aggregation and disaggregation of RBCs can help us to differentiate between the bridging and depletion mechanisms.

For a better characterization of differences between the bridging and depletion mechanisms, numerical simulations should be a suitable approach. Most of the existing models employ an attractive potential to induce aggregation between RBCs [22, 28, 35–37]. A combined experimental and simulation study [30] has shown that potential-based attractive interactions between RBCs reproduce well depletion-mediated aggregation. However, bridging interactions require explicit modeling of discrete bonds between cells. There exists a cross-linking model of RBC aggregation based on bond formation, which has been used to investigate the behavior of two-dimensional RBC doublets in shear flow [38]. In this model, temporal dynamics of the bonds is implemented through prescribed on- and off-rates for bond formation and dissociation. A similar modeling approach has to be employed to describe bridging interactions between RBCs. Furthermore, existing experimental observations [33, 34] suggest that bridges between RBCs might be mobile, such that they can slide along the RBC membrane without breakage. This can occur due to a non-specific adsorption of the bridging macromolecules to the RBC membrane or through the mobility of intra-membrane structures which interact with the bridging molecules [39].

In this work, we investigate dynamic aggregation and disaggregation of two RBCs using three different models for aggregation interactions. The first model is based on an attractive potential, which mimics depletion interactions between the two cells. The second model assumes dynamic formation/dissociation of *immobile* bonds, which bridge two RBCs whenever the distance between their membranes is small enough. Finally, the third model is newly proposed here, and based on dynamic formation/dissociation of *mobile* bonds, whose end points can slide along the membrane surface. Our simulations show that dynamic aggregation of two RBCs is primarily due to depletion interactions, while RBC disaggregation has both depletion and bridging contributions. The bridging model with immobile bonds shows an abrupt separation of two RBCs during disaggregation, when the pulling force is large enough to break all formed bonds nearly at the same time. The disaggregation process in corresponding experiments does not exhibit an abrupt force drop, which strongly favors the bridging model with mobile bonds. Simulated force profiles of RBC disaggregation agree well with the corresponding experimental measurements, and help us differentiate between the bridging and depletion mechanisms. Similar models can also be used to better understand aggregation and adhesion between other biological cells.

## II. METHODS & MODELS

### A. Disaggregation of RBCs using optical tweezers

#### 1. Blood sample preparation

Blood was taken from a clinically healthy donor from the ulnar vein and collected into the tubes with ethylenediaminetetraacetic acid (EDTA) as an anticoagulant. To obtain blood plasma without platelets, the blood is centrifuged using Eppendorf MiniSpin (Eppendorf, Germany) for 10 minutes at room temperature at 3000 × *g*. The supernatant is then collected and centrifuged once more for 10 minutes at 6000× *g* at room temperature. After the centrifugation, the obtained platelet poor plasma is carefully collected. We investigate the disaggregation of RBC doublets in plasma and in several mixtures of plasma and phosphate-buffered saline (PBS). RBCs separated from the whole blood are diluted to 1 : 500 in plasma or in plasma mixed with PBS (Gibco) at volume ratios 9 : 1, 8 : 2, and 7 : 3 (plasma : PBS). A stock solution of polystyrene beads (3 *μm* in diameter with a standard deviation ≤ 0.1 *μm* by Fluka, Germany) is prepared by 1 : 100 dilution of their stock solution in PBS. Then, the obtained dilute solution of beads is added to the suspension of RBCs at volume ratio 1 : 10 (volume of beads buffer : volume of RBC suspension).

### 2. Laser tweezers setup

The lay-out of the experimental setup of laser tweezers used in experiments is described Fig. 1(a). The setup is based on the inverted microscope (Eclipse TE 2000, Nikon, Tokyo, Japan), with multiple movable optical traps created using a laser beam generated by a single-mode Nd:YAG laser (1064 nm, Ventus, Laser Quantum, Stockport, United Kingdom). The beam is reflected by a parallelly-aligned liquid-crystal spatial light modulator (PAL-SLM, PPM X8267-15, Hamamatsu Photonics, Hamamatsu, Japan), and then focused through a large numerical aperture oil immersion objective (NA 1.25, 60x, Nikon, Tokyo, Japan). The position of the traps within the focal plane of the objective is controlled by the PAL-SLM, with adjustments made through software in the MATLAB environment. Visualization of the trapped cells is achieved using a digital CMOS camera (Orca-Flash 4.0 V3, Hamamatsu Photonics, Hamamatsu, Japan).

**FIG. 1.**
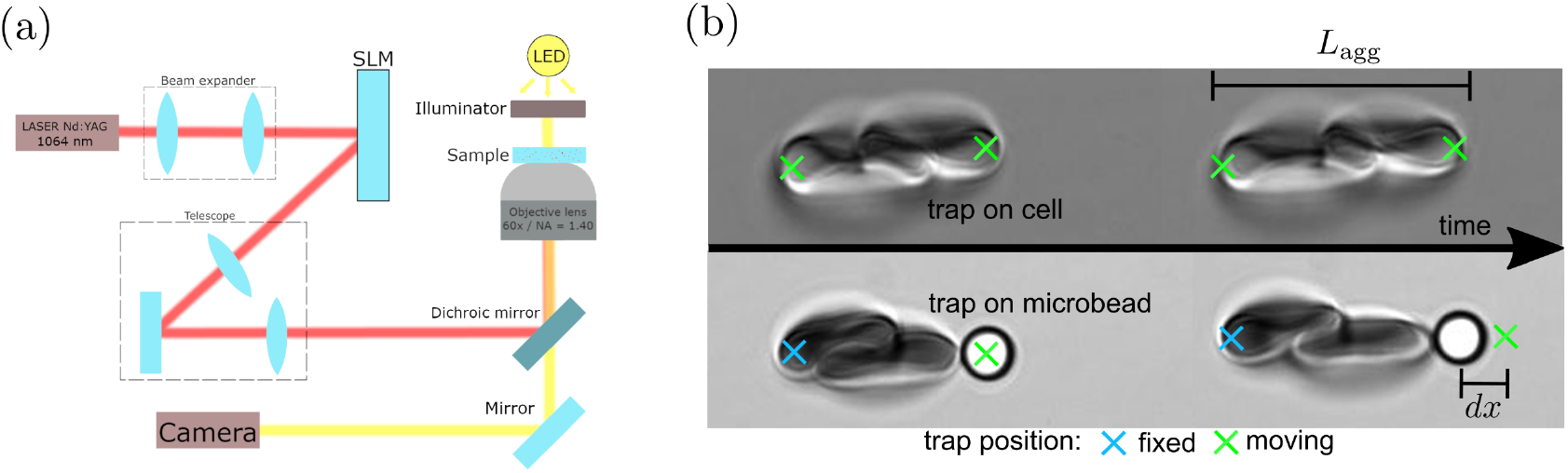
(a) Graphical lay-out of the laser tweezers setup. (b) Illustration of the experimental approach to measure the length *L*_*agg*_ of the doublets (top) and the disaggregation force between RBCs (bottom).

In our experiments, we consider two methods for the separation of aggregated RBCs using optical tweezers, as depicted in Fig. 1(b). The experiment starts by trapping two separate RBCs, where each cell is held by two optical traps, one on each RBC edge. Next, we bring the two cells face-to-face together, and deactivate the two traps located in the middle of the doublet to facilitate their aggregation. After the confirmation of stable RBC aggregation, we consider two strategies for pulling them apart: (i) *direct pulling*, where two RBCs are directly pulled away from each other using optical traps [see the top picture in Fig. 1(b)], and (ii) *bead-assisted pulling*, where an attached polystyrene bead is pulled away using the optical trap [see the bottom image in Fig. 1(b)]. The bead is attached to RBC membrane prior the doublet formation by putting the trapped bead in contact with a RBC for a few seconds until a firm adhesion is established. For the disaggregation phase, only two traps are used, including a stationary trap on the left side of one aggregated RBC [marked by a blue cross in Fig. 1(b)] and a movable trap on the right side of the other RBC. The movable trap is indicated by a green cross in Fig. 1(b), and holds the RBC on the right either directly or through the attached microbead. The disaggregation of RBCs proceeds by displacing the movable trap to the right.

### 3. Experimental measurement of the disaggregation force

Bead-assisted disaggregation was used to measure the interaction force between two RBCs during their disaggregation (i.e. disaggregation force). Exact measurements of the RBC disaggregation force are only available utilizing trapping of the microbead, since RBC is deformed upon trapping and its shape is significantly different from the circular, which complicates the tweezers calibration. Calibration of the laser tweezers setup is crucial for accurate determination of the trapping force exerted by the optical tweezers on the trapped object as a function of the laser’s power. We employ the calibration method based on Brownian motion, described in detail in Ref. [40]. This method involves the observation of the displacement of a microbead trapped by the laser tweezers through direct microscopy. The key steps of this calibration process are outlined in Ref. [41]. In the experiments, the disaggregation force is measured using the bead-assisted pulling scheme. Throughout the disaggregation process, we continuously track the positions of both the trapped bead and the optical tweezers. Initially, we align the center of the bead with the position of the optical tweezers. However, as the experiment progresses, the disaggregation force causes a misalignment between the bead’s center and the tweezers position. By measuring the offset (*dx*) between the trap’s position and the bead’s center, we compute the disaggregation force at any given moment as

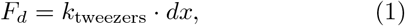

where *k*_tweezers_ is the stiffness of the trap. All the measurements are conducted at room temperature (293 K) using a laser power of 473 mW. The stiffness of the tweezers is *k*_tweezers_ = 27.2 pN*/μ*m based on the calibration described in Ref. [41]. The use of optical traps using PAL-SLM imposes certain limitations, notably that the right trap cannot be moved continuously. Instead, it is displaced in a step-wise manner, allowing the system to relax during each step. The displacement magnitude *dx*_step_ of the moving trap is set to 0.2 *μm*, and the interval between steps is *t*_*relax*_ = 3 seconds. These values correspond to the minimal displacement and relaxation time in our experimental setup. We construct the experimental disaggregation traces by measuring the end-to-end distance *L*_*agg*_ of the doublets and *F*_*d*_.

### B. Red blood cell model

A RBC membrane is represented by a triangulated network model [see Fig. 2(a)], which properly accounts for shear elasticity and bending rigidity of the cells [42, 43]. This model has been successfully used in simulations of RBC aggregation [28] and blood flow [44]. Each RBC is modelled by a triangulated network of *N* = 3000 vertices and *N*_*s*_ = 3(*N*− 2) springs, see Fig. 2(a). The total potential energy of the membrane is a sum of in-plane elastic energy *U*_elastic_, bending energy *U*_bend_, and surface-area and volume constraints (*U*_area_ and *U*_vol_) as *U*_total_ = *U*_elastic_ + *U*_bend_ + *U*_area_ + *U*_vol_. The elastic energy is given by

**FIG. 2.**
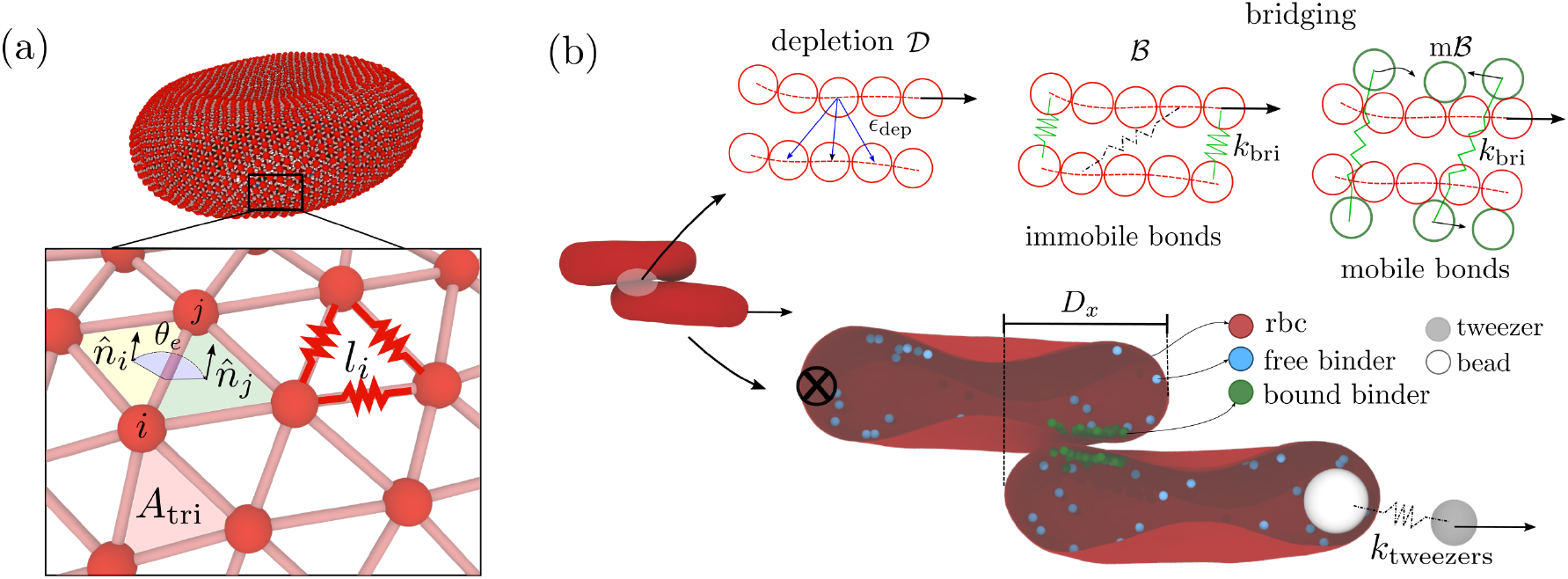
(a) Triangulated membrane model of a RBC. The membrane is represented by a set of vertices connected by springs. (b) Schematic representation of aggregation interactions (depletion and bridging) between two RBCs and the setup for modelling disaggregation dynamics mimicking optical tweezers.

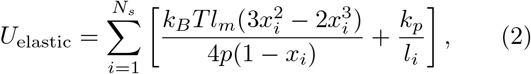

where the first term is an attractive worm-like-chain potential with the persistent length *p*, the length *l*_*i*_ of the *i*-th spring, the maximum spring extension *l*_*m*_, and *x*_*i*_ = *l*_*i*_*/l*_*m*_, while the second term is a repulsive potential with the coefficient *k*_*p*_. Force balance between the attractive and repulsive potential yields a non-zero equilibrium spring length.

The bending energy is given by

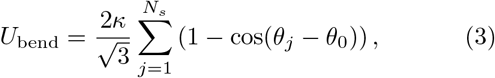

where *κ* is the bending modulus from the Helfrich curvature-elasticity model [45], *θ*_*j*_ is the angle between the normals of the two triangles sharing the *j*-th edge, and *θ*_0_ is the spontaneous angle. The surface-area constraint is given by

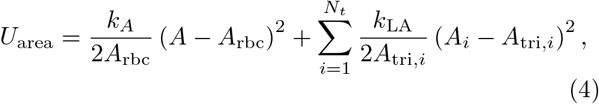

where *A* is the total area of the membrane, *A*_rbc_ is the target area of the cell, *A*_*i*_ is the area of the *i*-th triangle, *A*_tri,*i*_ is the target area of the *i*-th triangle, *k*_*A*_ is the total-area constraint coefficient, and *k*_LA_ is the local-area constraint coefficient. Finally, the volume constraint is expressed as

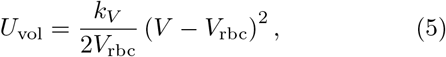

where *V* is the RBC volume, *V*_rbc_ is the target volume of the RBC, and *k*_*V*_ is the volume constraint coefficient.

### C. Aggregation models

Following the two hypotheses (i.e., depletion and bridging) for RBC aggregation, we consider three different models: (i) 𝒟 *model* with the aggregation mediated by an attractive interaction potential, mimicking depletion interactions, (ii) ℬ *model* that bridges the two cells by non-mobile bonds, and (iii) *m* ℬ *model* where RBC-RBC interactions are implemented through mobile bonds [see Fig. 2(b)]. The depletion model (𝒟) implements RBC aggregation through a truncated Lennard-Jones (LJ) potential given by

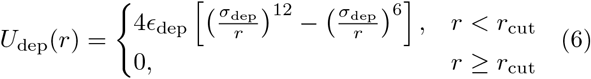

where *ϵ*_dep_ is the potential depth, *σ*_dep_ is the characteristic repulsion length, and *r*_cut_ is the cutoff radius beyond which the potential vanishes. The LJ potential is applied to every pair of vertices belonging to distinct RBCs. A similar interaction potential has already been used to impose an effective non-specific aggregation between RBCs [24, 25, 28, 46]. However, a recent study [30] has shown that an interaction potential due to depletion induced by suspended *fd* -virus rod-like particles is similar to a potential imposed by the attractive LJ interactions, where the interaction strength *ϵ*_dep_ is proportional to the concentration of suspended rod-like particles. Note that the total adhesion energy between two RBCs is a function of the number of interacting vertices (i.e., contact area) and the interaction strength *ϵ*_dep_ [28].

Bridging is implemented through a dynamic formation and dissociation of bonds between the two RBCs. A bond is formed when a distance between two not-bound bridge-forming particles is smaller than *r*_form_, while an existing bond dissociates when its length is larger than *r*_break_. The bonds are modelled by a harmonic potential

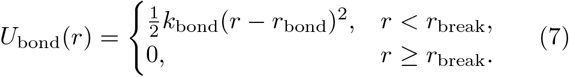

where *k*_bond_ and *r*_bond_ are the bond strength and equilibrium distance, respectively. Note that a more sophisticated bond formation/dissociation model (e.g., with a stress-dependent off-rate) can easily be considered. However, we expect that more sophisticated bond models would lead to qualitatively similar dynamics of RBC disaggregation in comparison to the employed simple model, because the relaxation time *t*_*relax*_ between consecutive displacements of the optical tweezers is significantly larger than characteristic times for bond formation and dissociation.

For the ℬmodel, bonds can form between every not-bound pair of vertices belonging to different RBCs, see Fig. 2(b). In this case, the bonds are not mobile, as they connect different RBC vertices, which are part of an elastic network. The surface density of particles that are able to form bonds is simply determined by the RBC resolution as *ρ*_sites_ = *N/A*_rbc_. For the *m*ℬ model, bond-forming particles should become mobile. Since it is difficult to introduce the surface mobility of RBC vertices directly, we place *n*_*b*_ mobile particles inside each RBC, which represent potential bond-forming partners, see Fig. 2(b). To prevent crossing of the membrane by the bond-forming particles inside the RBCs, the repulsive part of the LJ potential in Eq. (6) is used with *r*_cut_ = 2^1*/*6^*σ*_dep_. The surface density of these internal binders for the *m* model is then *ρ*_sites_ = *n*_*b*_*/A*_rbc_. Note that binders can freely move inside the RBCs, and only binder pairs belonging to distinct cells can form bonds. Excluded-volume interactions between binders within the same RBC are also implemented through the repulsive part of the LJ potential in Eq. (6).

### D. Simulation setup and parameters

To mimic the experimental setup on aggregation and disaggregation of RBCs using optical tweezers [see Fig. 1(b)], we place two RBCs close enough to each other to initiate aggregation, with different overlapping distances *D*_*x*_. One of the cells is immobilized by fixing a fraction (70 particles) of RBC vertices on its left side, as shown in Fig. 2(b) by the black cross. In our optical tweezers experiments, the applied strain is controlled, such that step-wise displacements are imposed on the tweezers position and the force exerted on the bead is measured. To mimic the experimental setup, we place a bead *B*_*i*_ with a radius of 0.9 *μm* inside the moving RBC, drawn in white in Fig. 2(b). The bead *B*_*i*_ is attached to another bead *B*_*o*_ outside the RBC [grey circle in Fig. 2(b)] by a harmonic spring with a coefficient *k*_tweezers_ and an equilibrium distance *r*_*B*_. To mimic the dynamic operation of optical tweezers, the bead *B*_*o*_ is pulled using step-wise displacements of 0.2 *μ*m with a relaxation time of 0.5 s between consecutive steps. The applied force *F*_*d*_ is monitored through the extension *r* of the spring as *F*_*d*_ = *k*_tweezers_(*r*− *r*_*B*_). The bead *B*_*i*_ stays inside the moving RBC due to purely repulsive LJ interactions between *B*_*i*_ and RBC vertices.

Evolution of the simulation system follows a Langevin dynamics equation given by

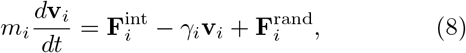

where *m*_*i*_ is the mass of particle *i* (including RBC vertices, *B*_*i*_, *B*_*o*_, and binder particles if used), **v**_*i*_ is the particle velocity,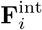 is the total conservative force on particle *i* due interactions with other particles, *γ*_*i*_ is the friction coefficient, and 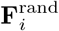 is the random force accounting for thermal fluctuations. Following a fluctuation-dissipation relation, the random force satisfies 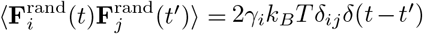. Note that Langevin dynamics models effectively the dissipation from surrounding fluids, but does not provide hydrodynamic interactions.

Table I presents main simulation parameters in both model and physical units. We define a length scale through an effective RBC diameter 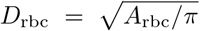 with the membrane area *A*_rbc_ = 133 in simulations, corresponding to *A*_rbc_ = 133.5 *μ*m^2^. Furthermore, the energy unit is *k*_*B*_*T* (*k*_*B*_*T* = 0.1 in simulations) for the ambient temperature of 37^*°*^ C, and a time scale is defined as *τ* = *ηD*_rbc_*/Y*, where *η* is the dynamic viscosity of a suspending fluid, and *Y* is the Young’s modulus of RBC membrane. Even though we do not simulate the surrounding fluid explicitly, we use *η* to compute the friction coefficient on RBC particles as *γ*_*rbc*_ = 3*πηD*_rbc_*/N* and on other particles (e.g., *B*_*i*_, *B*_*o*_, and binder particles) as *γ*_*p*_ = 6*πηR*_*p*_ with a radius *R*_*p*_. The dynamic viscosity in physical units is assumed to be *η* = 1.2 × 10^*−*3^ Pa· s. The average Young’s modulus of a healthy RBC is *Y* = 18.9 *μ*Nm^*−*1^ [43]. The mass for all particles in simulations is set to *m* = 1. The simulation time step is Δ*t* = 0.003*τ*. Unless stated otherwise, for the ℬmodel, we employ *r*_bond_ = 0.046*D*_rbc_, *r*_form_ = 0.077*D*_rbc_, and *r*_break_ = 0.108*D*_rbc_, while for the *m*ℬ model, *r*_bond_ = 0.077*D*_rbc_, *r*_form_ = 0.108*D*_rbc_, and *r*_break_ = 0.154*D*_rbc_ are set to take into account a finite thickness of the RBC membrane.

Different parameters during disaggregation of two RBCs are analyzed, including the overlapping distance *D*_*x*_, end-to-end distance *L*_*agg*_ of the doublet, the separation *d*_*cc*_ of two centers of mass, RBC deformation, and the disaggregation force *F*_*d*_. RBC deformation is analyzed through the eigen-values *I*_1_ *> I*_2_ *> I*_3_ of the gyration tensor *G*_*ij*_ computed as

**TABLE I.** Simulation parameters in model and physical units. RBC parameters in physical units correspond to average properties of healthy RBCs. In all simulations, *A*_rbc_ = 133, *k*_*B*_*T* = 0.1, and *η* = 97.8 were set.

| Parameters | Model units | Physical units |
| --- | --- | --- |
| number of vertices $N$ | 3000 | |
| membrane area $A_{\text{rbc}}$ | 133 | $133.5 \mu\text{m}^2$ |
| energy unit $k_B T$ | 0.1 | $4.28 \times 10^{-21} \text{ J}$ |
| fluid viscosity $\eta$ | 97.8 | $1.2 \times 10^{-3} \text{ Pa} \cdot \text{s}$ |
| RBC diameter $D_{\text{rbc}}$ | $\sqrt{A_{\text{rbc}}/\pi}$ | $6.5 \mu\text{m}$ |
| RBC volume $V_{\text{rbc}}$ | $0.34 D_{\text{rbc}}^3$ | $93 \mu\text{m}^3$ |
| Young's modulus $Y$ | $1.82 \times 10^5 k_B T / D_{\text{rbc}}^2$ | $18.9 \mu\text{Nm}^{-1}$ |
| bending modulus $\kappa$ | $70 k_B T$ | $3 \times 10^{-19} \text{ J}$ |
| local area coefficient $k_{\text{LA}}$ | $4.2 \times 10^4 k_B T / D_{\text{rbc}}^2$ | $4.3 \mu\text{Nm}^{-1}$ |
| total area coefficient $k_A$ | $2.1 \times 10^6 k_B T / D_{\text{rbc}}^2$ | $2.1 \times 10^2 \mu\text{Nm}^{-1}$ |
| volume coefficient $k_V$ | $1.4 \times 10^7 k_B T / D_{\text{rbc}}^3$ | $220 \text{ Nm}^{-2}$ |
| time scale $\tau$ | $\eta D_{\text{rbc}} / Y$ | $4.1 \times 10^{-4} \text{ s}$ |
| tweezers spring coefficient $k_{\text{tweezers}}$ | $5.33 \times 10^4 k_B T / D_{\text{rbc}}^2$ | $5.4 \mu\text{Nm}^{-1}$ |
| tweezers displacement step $dx_{\text{step}}$ | $0.031 D_{\text{rbc}}$ | $0.2 \mu\text{m}$ |
| relaxation time $t_{\text{step}}$ | $1210 \tau$ | $0.5 \text{ s}$ |

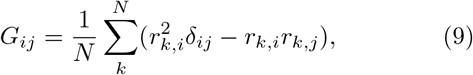

where *i, j* ∈ { *x, y, z* }, and (*r*_*k,x*_, *r*_*k,y*_, *r*_*k,z*_) is the position vector of the *k*-th particle with respect to the RBC center of mass.

## III. RESULTS

### A. Spontaneous RBC aggregation

Aggregation of two RBCs or vesicles has been studied both in experiments [27, 30] and simulations [28, 47, 48], predicting a variety of doublet conformations as a function of depletion or interaction strength. Here, we demonstrate that spontaneous aggregation of two RBCs occurs when they are close enough to each other, as it is observed experimentally [27, 30]. We employ the 𝒟 model that mimics depletion interactions with the attraction strength in the range *ϵ*_dep_ = 0.1*k*_*B*_*T* − 1.0*k*_*B*_*T*. This range of *ϵ*_dep_ represents a moderate aggregation strength, such that aggregated RBCs do not visually deform much, which is consistent with RBC aggregation in plasma. Here, we employ *σ*_dep_ = 0.031*D*_rbc_ and *r*_cut_ = 2.5*σ*_dep_ for the LJ potential in Eq. (6). Starting from a minimal contact of two cells, they aggregate by maximizing their contact area. Eventually, the two RBCs attain a stacked configuration with a maximum contact area. The aggregation time depends on the initial overlap and the strength of attractive interactions.

Spontaneous RBC aggregation through the formation of bonds is different from that due to the depletion interactions. For the ℬ model, when two RBCs are brought close to each other, bonds are first formed within the initial contact area. However, starting from a small initial overlap of the RBCs, we do not observe their sliding along each other to maximize the contact area, as it was observed for the depletion interactions with the 𝒟 aggregation model. As the binding strength mediated by *k*_bond_ is increased, more contact area between the two RBCs can be gained through zipping of the two membranes at the contact edges, while membrane sliding relative to each other is prohibited due to the immobility of already formed bonds. For the *m* ℬ model with mobile bonds, the relative sliding of membranes does occur, although the spontaneous aggregation process is much slower than for the 𝒟 model, because the sliding motion is limited by the mobility of free binders. As a result, our simulations suggest that spontaneous RBC aggregation with an increase in contact area, which is also observed experimentally, is primarily due to depletion interactions, while bridging (if present) plays at most a secondary role in this process. Therefore, the ℬ and *m* ℬ models are employed further in combination with the 𝒟 model, which is also presumably the case in experiments on RBC aggregation due to the presence of various suspended macromolecules.

### B. Disaggregation of two RBCs

Several experimental studies [31, 32] report that the force required for RBC doublet disaggregation increases with increasing time of contact, and saturates at large enough times. Clearly, pure depletion interactions cannot account for this time dependence, because the disaggregation force in this case should depend only on the concentration of depletant and the contact area between the two cells. It is hypothesized that this contact-time dependence of the disaggregation force is due to a characteristic time of the formation of bonds (or bridges) [32]. For the ℬmodel with immobile bonds, the total number of bonds depends only on the contact area between the cells, whereas for the *m*ℬ model, the total number of bonds can be affected by the time of contact, since the binders diffuse inside the RBCs and participate in bond formation only when they are close enough to a free binding partner. To eliminate the effect of contact time in the comparison of different aggregation mechanisms in our simulations, the two RBCs are let initially to aggregate for about 2 s, before the disaggregation process is started. We have verified that the initial contact time of 2 s is long enough to achieve full saturation of the aggregation interactions for the investigated ranges of simulation parameters (see Ref. [41]).

#### 1. Disaggregation of RBCs with depletion interactions

Using the 𝒟model, we investigate the effect of pure depletion interactions on the disaggregation of two RBCs. Figure 3(a) shows *F*_*d*_ as a function of the overlap distance *D*_*x*_ for several different initial-overlap distances. Starting from a non-zero initial overlap (i.e., larger *D*_*x*_ values), *F*_*d*_ has first a steep increase, because the applied force must exceed drag and aggregation forces before the pulled RBC starts sliding with respect to the held cell. Furthermore, the initial increase in *F*_*d*_ is also partially associated with RBC deformation that stores some amount of elastic energy. When a critical value of *F*_*d*_ is reached, the pulled RBC starts moving with a nearly constant velocity, and the disaggregation force decreases with decreasing overlap distance *D*_*x*_. This decay in *F*_*d*_ is expected, because depletion forces are proportional to the contact area, which decreases when the two RBCs are pulled apart. Consistently, the maximum in *F*_*d*_ increases with increasing initial overlap or the contact area. However, the decay in *F*_*d*_ becomes independent of the initial overlap of the cells, after a critical force for the disaggregation is reached. Note that right before RBC separation (*D*_*x*_ ≈0), there is a slight increase (≤ 1.5 pN) in the disaggregation force. This occurs due to a change in the alignment of the two RBCs right before the separation, which is accompanied by a small increase in their contact area, see Fig. 3(c). After the cells are fully separated (*D*_*x*_ *<* 0), *F*_*d*_ is equal to the drag force exerted by the fluid on the pulled RBC with *F*_drag_ ∼ 5 pN. Therefore, depletion forces between the two RBCs can be computed as *F*_*d*_− *F*_drag_, and are in the range of 3−5 pN, as shown in Fig. 3(a).

**FIG. 3.**
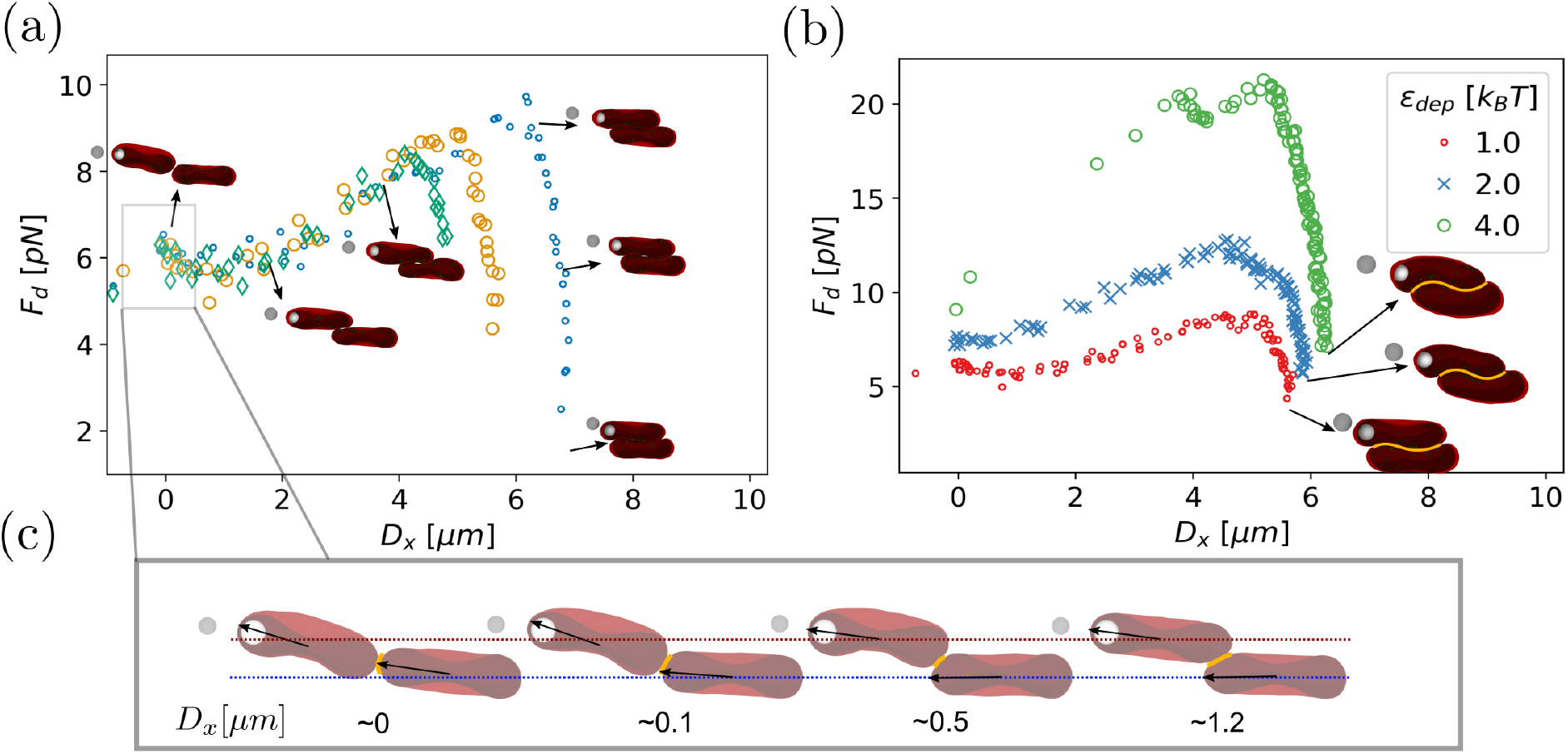
Disaggregation force for the depletion-mediated model. (a) *F*_*d*_ as a function of *D*_*x*_ for different initial overlap distances of two RBCs at a fixed depletion strength of *ϵ*_dep_ = 1*k*_*B*_*T*. (b) *F*_*d*_ dependence for different depletion strengths *ϵ*_dep_, starting from the initial overlap of 60%. (c) Illustration of final separation of the two cells, indicating the approximated overlapping distance *D*_*x*_.

Figure 3(b) presents the disaggregation force *F*_*d*_ for different attraction strengths *ϵ*_dep_, starting from a fixed initial overlap of ≈ 60%. As *ϵ*_dep_ increases, significantly larger disaggregation forces are required to separate the two RBCs, as well as the induced RBC deformation is much more pronounced. Nevertheless, the dependence of *F*_*d*_ on *D*_*x*_ is qualitatively similar for various *ϵ*_dep_ values. After the critical disaggregation force is reached for the case of *ϵ*_dep_ = 4*k*_*B*_*T*, a non-monotonic decrease in *F*_*d*_ is due to energetic costs associated with doublet deformation during disaggregation.

#### 2. Disaggregation force from immobile bonds

Figure 4(a,b) shows disaggregation forces for two different initial overlap distances (60% and 100%) and various strengths *k*_bond_ of bonds. As already mentioned, the ℬ model is used in combination with the 𝒟model (*ϵ*_dep_ = 1*k*_*B*_*T*), which aids in the initial aggregation of the two RBCs. Similarly to the 𝒟model, the model with immobile bridges shows a steep increase in *F*_*d*_ after starting the disaggregation process. As expected, the maximum disaggregation force is proportional to the strength of bonds and the initial overlap area between the cells, because the number of initially formed bridges is proportional to the contact area. However, shortly after the maximum in *F*_*d*_ is reached, a sudden reduction in the force occurs, which is qualitatively different from a relatively slow decay in *F*_*d*_ for the 𝒟 model mimicking depletion interactions. The abrupt decay in *F*_*d*_ for the ℬmodel is associated with the breakage of nearly all bridges within a short time due to bond stretching, which results in a sudden reduction in the contact area between the two RBCs. Note that for all cases in Fig. 4(a,b), final force after the separation is *F*_*d*_ ≈ *F*_drag_ as expected. However, the separation of two RBCs is often attained when they still overlap (i.e., for *D*_*x*_ *>* 0), indicating that the breakage of all bonds occurs earlier than they reach a side-to-side configuration with *D*_*x*_≈ 0. The disaggregation force is larger for the case of 100% initial overlap with *D*_*x*_ = 8 *μ*m in Fig. 4(b) than for the 60% initial overlap in Fig. 4(a), due to a larger number of formed bonds.

**FIG. 4.**
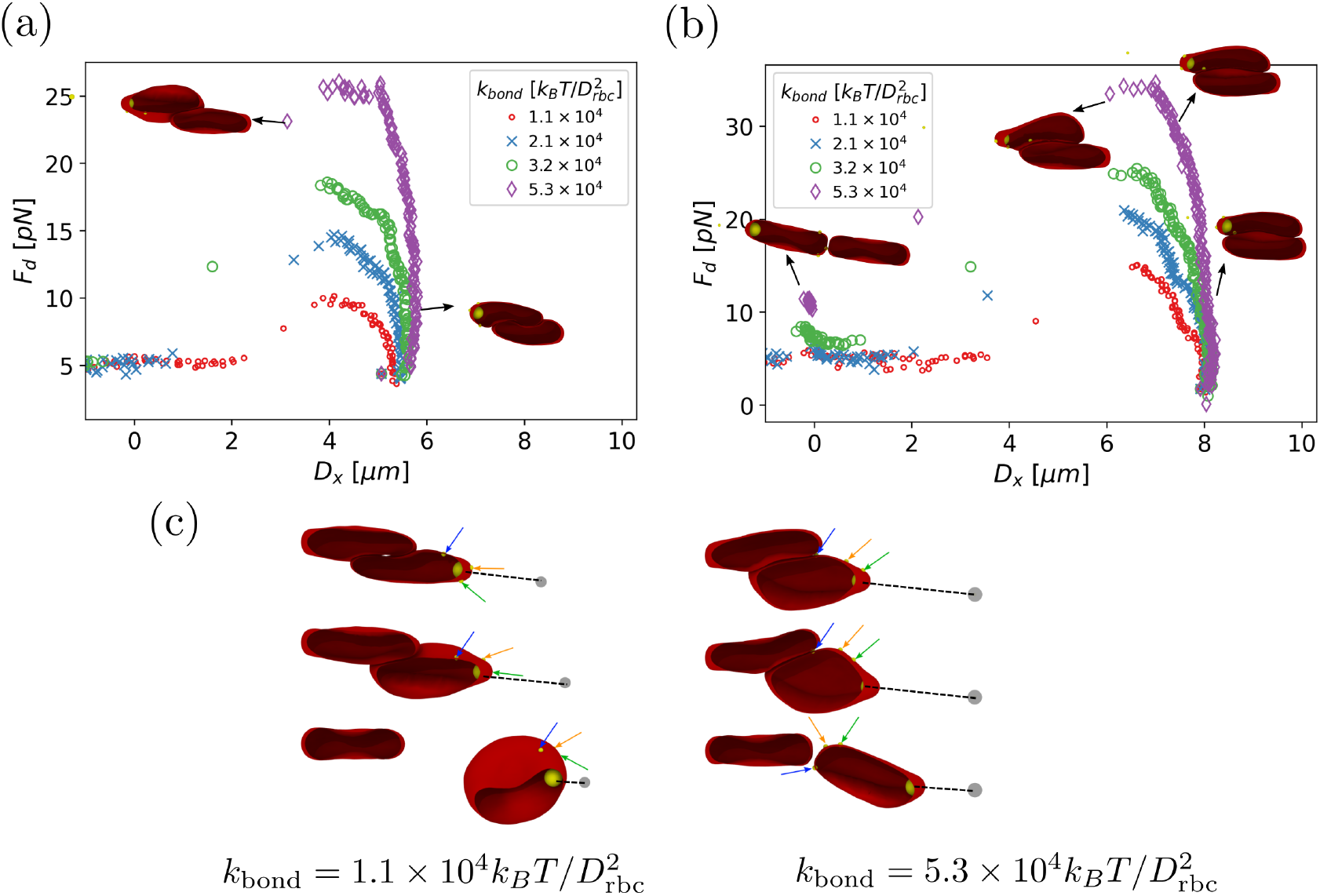
Disaggregation force from the ℬ model for different bond strengths. Two initial overlaps, including (a) 60% with *D*_*x*_ = 5 *μ*m and (b) 100% with *D*_*x*_ = 8 *μ*m, are considered. (c) Illustration of a tank-treading-like motion of the RBC membrane for the *B* model with 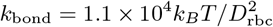 and 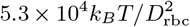. The arrows indicate the position in time of three markers on the membrne surface. Membrane dynamics is enhanced with increasing strength of the formed bridges, and the separation of two RBCs is generally abrupt.

Disaggregation dynamics of the RBC membranes for the 𝒟and ℬmodels is also very different. For depletion interactions, the two membranes slide along each other. In contrast, for the case of immobile bonds, disaggregation resembles peeling off one membrane from the other, because lateral sliding is restricted by the bonds. As a result, the RBC membranes undergo a tank-treading-like motion, which is often observed for RBCs in shear flow [49–51]. Figure 4(c) presents a time sequence of several snapshots for two different bond strengths 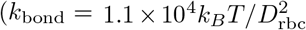 and 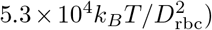, where three selected markers at the RBC membrane demonstrate its tank-treading-like motion. Furthermore, for the low bond strength with 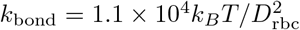, the detachment of the cells occurs abruptly when they still have a *D*_*x*_ *>* 0 overlap, see Fig. 4(c). In contrast, for the stronger bonds with 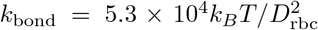, the tank-treading-like motion is more pronounced, and the separation of two cells occurs when the side-to-side configuration with *D*_*x*_ ≈ 0 is attained. Here, the tanktreading-like motion facilitates peeling off one membrane from the other by applied disaggregation forces.

#### 3. Disaggregation of RBCs connected by mobile bridges

Finally, we explore the disaggregation response of two RBCs, using the *m*ℬ model with mobile bonds. Here, the adhesion strength can be mediated by the density *ρ*_sites_ of binders and the bond strength *k*_bond_. Figure 5(a) shows *F*_*d*_ as a function of *D*_*x*_ for *ρ*_sites_ = 106*/A*_rbc_ and 1463*/A*_rbc_, and two strengths of depletion interactions *ϵ*_dep_ = 0.1*k*_*B*_*T* and *k*_*B*_*T*. As expected, *F*_*d*_ substantially increases when the density of binders becomes large, due to a large number of bonds that need to be broken for the separation of RBCs. For the low density of binders, the disaggregation process is governed at the beginning by depletion interactions, which can be seen in Fig. 5(a) through a nearly constant *F*_*d*_ ≈ 5 pN for *ϵ*_dep_ = 0.1*k*_*B*_*T* and a characteristic elevation in *F*_*d*_ for *ϵ*_dep_ = *k*_*B*_*T*. Note that *F*_*d*_ ≈ 5 pN corresponds to the drag force of a single RBC in our simulations. In the case of *ρ*_sites_ = 106*/A*_rbc_, the formed bonds between two RBCs are able to move along the membrane, leading to a nearly negligible contribution to *F*_*d*_ during the initial stage of the disaggregation process. However, when *D*_*x*_ →0, there is an increase in *F*_*d*_, which is due to the accumulation of several mobile bridges near the RBC edges that need to be eventually broken. For the case of *ρ*_sites_ = 1463*/A*_rbc_, the disaggregation force increases substantially at the beginning, which resembles the behavior of *F*_*d*_ for the ℬmodel with immobile bonds. Due to a significant crowding of formed bonds, their contribution to measured *F*_*d*_ is quite pronounced. However, an abrupt separation of RBCs, as observed for the ℬmodel, does not take place here due to the mobility of bonds. Note that for large *ρ*_sites_, there is no increase in *F*_*d*_ when *D*_*x*_ → 0.

**FIG. 5.**
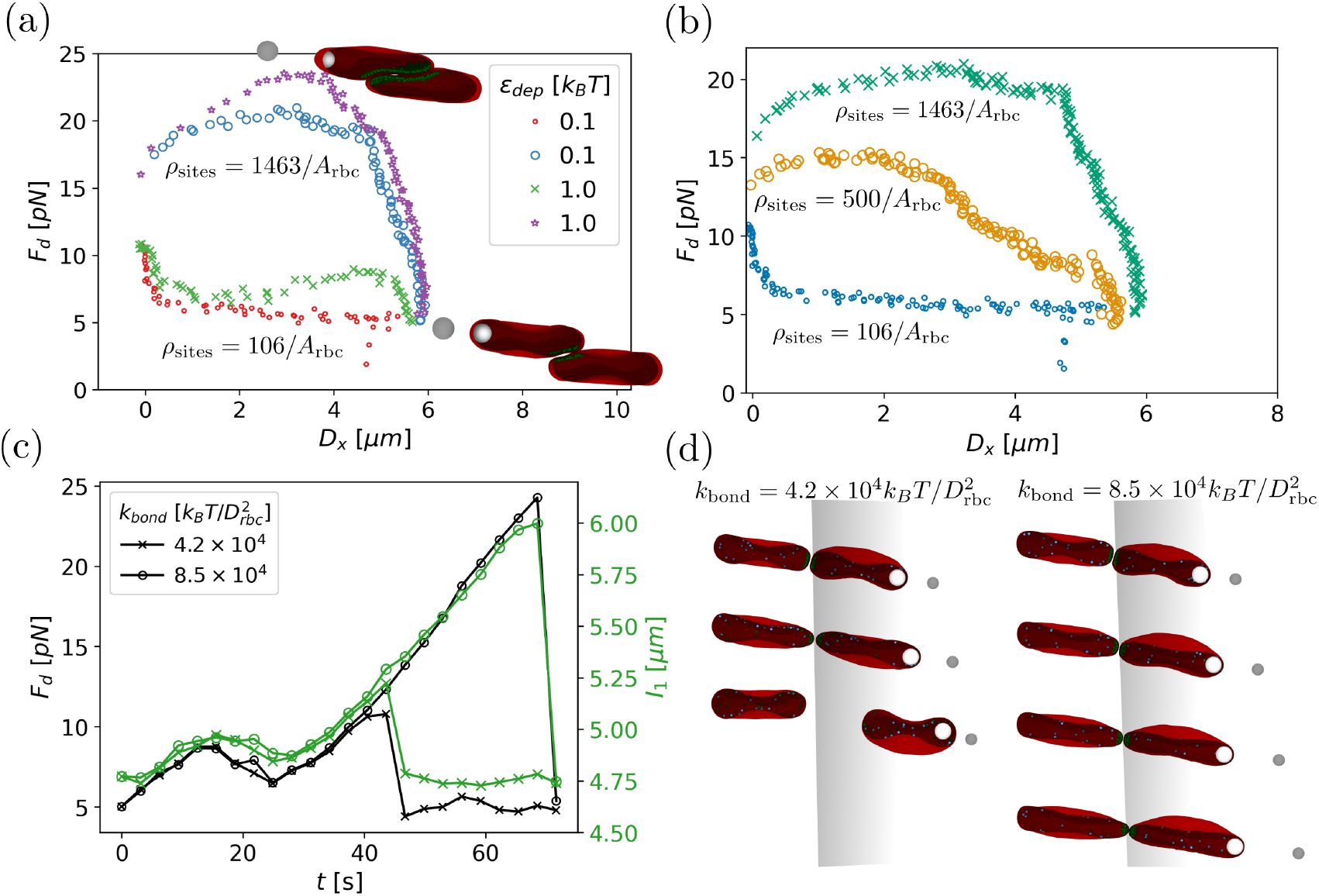
Disaggregation force for different surface densities *ρ*_sites_ and strengths *k*_bond_ of mobile bonds. (a) *F*_*d*_ for two depletion interaction strengths (*ϵ*_dep_ = *{*0.1, 1*}k*_*B*_*T*) and two surface densities (*ρ*_sites_ = *{*106, 1463*}/A*_rbc_) of mobile binders, with an initial overlap of 60% and 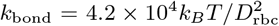. (b) Comparison of disaggregation forces for three different surface densities *ρ*_sites_ = *{*106, 500, 1463*}/A*_rbc_ without depletion interactions and 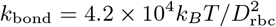. (c) Disaggregation force *F*_*d*_ and RBC elongation *I*_1_ for two different bond strengths. Here, *ρ*_sites_ = 106*/A*_rbc_. (d) Snapshots of the RBCs for two different bond strengths right when the detachment occurs.

To emphasize differences between the 𝒟and *m*ℬ models, we also perform simulations with an intermediate density *ρ*_sites_ = 500*/A*_rbc_ of binders. Figure 5(b) compares *F*_*d*_ profiles for different *ρ*_sites_ values without depletion interactions. For the case of *ρ*_sites_ = 500*/A*_rbc_, an initial increase in *F*_*d*_ at the beginning of the disaggregation process is similar to that due to depletion interactions, as in the case of *ρ*_sites_ = 106*/A*_rbc_ and *ϵ*_dep_ = *k*_*B*_*T* in Fig. 5(a). However, *F*_*d*_ continues to increase with decreasing *D*_*x*_ for the *m*ℬ model, while *F*_*d*_ decreases for the 𝒟model. Thus, the 𝒟and *m*ℬ models can be distinguished well for low and moderate strengths of aggregation.

Figure 5(c) presents the time evolution of the disaggregation force *F*_*d*_ and the RBC elongation characterized by the largest eigen-value *I*_1_ of the gyration tensor for different strengths of mobile bonds 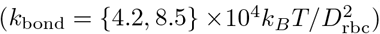 at *ρ*_sites_ = 106*/A*_rbc_. As expected, the elongation of RBCs is strongly correlated with the magnitude of *F*_*d*_. The bond strength *k*_bond_ does not significantly affect the initial behavior of *F*_*d*_, because of the dominance of depletion interactions at the beginning of the disaggregation process for low *ρ*_sites_, while the final disaggregation force is proportional to *k*_bond_. Regardless of the strength of interactions in the *m*ℬ model, the final breakage of bonds occurs at the side-to-side configuration with *D*_*x*_ ≈ 0. Figure 5(d) provides several snapshots of RBCs as they detach from each other. Clearly, the elongation of cells is more pronounced for stronger bonds (i.e., with a larger *k*_bond_), and the detachment occurs irreversibly without the possibility for new adhesive interactions between the two RBCs.

#### 4. Experiments using optical tweezers

Several experimental studies [27, 30, 32] have demonstrated spontaneous aggregation of RBCs in plasma or other depletion-inducing solutions. Interestingly, our experiments of RBC aggregation in mixtures with different plasma:PBS ratios show that the tendency for spontaneous aggregation reduces as the dilution of plasma with PBS increases. This is consistent with our simulation results, which suggest that spontaneous aggregation of RBCs is mediated by depletion interactions. Through the dilution by PBS, the concentration of macromolecules in the suspending solution decreases, which leads to weaker depletion interactions, and a reduced tendency for spontaneous aggregation of the two cells.

To test the plausibility of different aggregation models, we also conduct RBC disaggregation experiments in plasma and plasma/PBS mixtures using optical tweezers. Figure 6(a) presents the disaggregation force for RBCs suspended in plasma and in a 8:2 mixture of plasma:PBS as a function of time. At the beginning, *F*_*d*_ exhibits an increase followed by a slight decrease, which qualitatively resembles the *F*_*d*_ dependence obtained in simulations using the 𝒟model mimicking depletion interactions. Then, there is a second increase in *F*_*d*_, which has been observed in simulations using the *m*ℬ model with mobile bonds. To approximate experimental measurements by simulations, we employ the combination of 𝒟and *m*ℬ models, as shown in Fig. 6(a). In the comparison, the disaggregation force is normalized by *k*_tweezers_*dx*_step_, which is the force exerted by the tweezers during a single displacement, while the time is normalized by the relaxation time *t*_step_ after each displacement. The simulated *F*_*d*_ curves follow well the experimental data, suggesting that both depletion and bridging interactions contribute to RBC aggregation. Evolution of the end-to-end distance *L*_agg_ of the RBC doublet is shown in Fig. 6(b), with a semiquantitative agreement between experimental and simulation measurements. Note that experimental disaggregation traces often exhibit variations in their absolute values, which can be attributed to differences in RBC properties and their aggregation interactions. To better illustrate the disaggregation process, Fig. 6(c) compares side-by-side experimental observations and simulation snapshots (see also Movie S1 [41]).

**FIG. 6.**
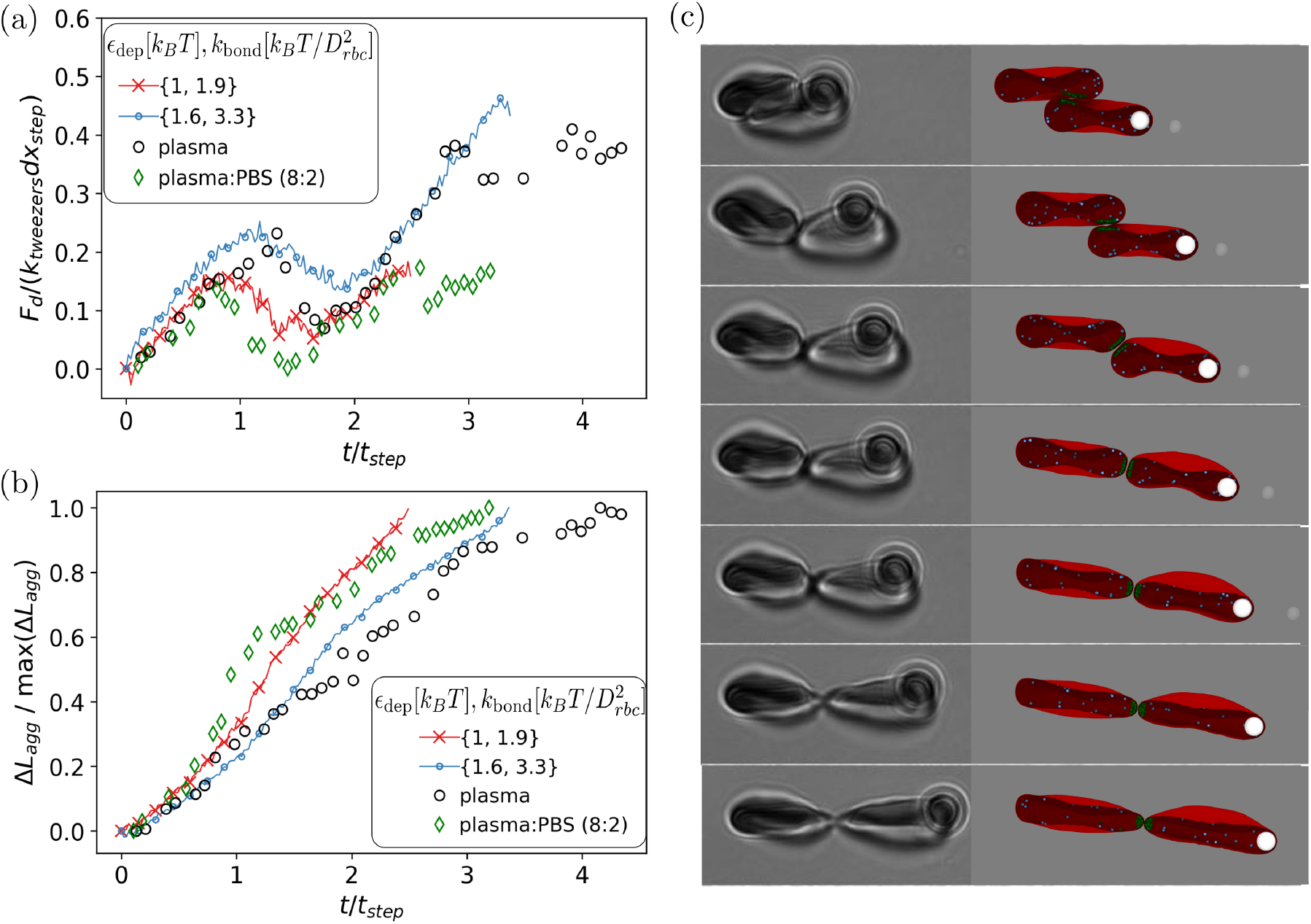
Comparison of simulated and experimental disaggregation of two RBCs. (a) Disaggregation force as a function of time. (b) End-to-end distance of a doublet during the disaggregation process. (c) Side-by-side comparison of experimental observations and simulation snapshots. Experiments are performed in pure plasma and in a 8:2 mixture of plasma:PBS. Simulations employ the *mB* model with additional depletion interactions. See also Movie S1 [41].

## IV. DISCUSSION AND CONCLUSIONS

We have systematically investigated the aggregation and disaggregation of two RBCs using three different models. For depletion interactions using the 𝒟model, the disaggregation force is proportional to the strength *ϵ*_dep_ of the attractive potential and the initial overlap area between RBCs. For bridging with immobile bonds (i.e., the ℬmodel), the maximum disaggregation force is proportional to the bond strength *k*_bond_ and the initial overlap area, since it determines the total number of formed bonds between cells. For bridging with mobile bonds (i.e., the *m*ℬ model), the maximum disaggregation force is proportional to the surface density *ρ*_sites_ of binders and the strength *k*_bond_ of the mobile bonds. For the 𝒟 and ℬ models, *F*_*d*_ exhibits an initial increase reaching a maximum force, followed by a continuous decrease in *F*_*d*_ for the 𝒟model or an abrupt drop in *F*_*d*_ for the ℬ model. In contrast, the *m* model yields two disaggregation regimes, depending on the binder density. At low densities of binders, an increase in *F*_*d*_ typically occurs when *D*_*x*_ → 0 (i.e., close to the time point of RBC separation), whereas at large *ρ*_sites_, *F*_*d*_ first increases and then decreases due to substantial crowding of the bonds. The mobility of bonds leads to a continuous decay in *F*_*d*_ in comparison with the abrupt drop in *F*_*d*_ for the ℬmodel with immobile bonds.

Figure 7(a) compares disaggregation forces for the three models as a function of *D*_*x*_ (see also Movies S2-S4 [41]). Here, the 𝒟 model assumes *ϵ*_dep_ = 1*k*_*B*_*T*, the ℬ model adopts 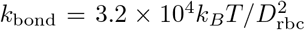, and the *m*ℬ model is based on 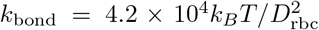 and *ρ*_sites_ = 106*/A*_rbc_, producing a comparable maximum disaggregation force of *F*_*d*_ ≈ 10 pN. RBC extension shown in Fig. 7(b) strongly correlates with *F*_*d*_, though the RBC elongation is slightly larger for the *m*ℬ model than for the model. This can also be seen from several snapshots at different time points in Fig. 7(c) for the three aggregation models. From these results, two different stages in the disaggregation process can be identified, as shown in Fig. 7(d-f). The first stage represents initial force yielding with nearly no change in *D*_*x*_, which can be mediated by both depletion and bridging interactions. In the second stage, the pulled RBC starts moving, where drag forces in addition to the depletion and bridging interactions contribute to the measurement of *F*_*d*_.

**FIG. 7.**
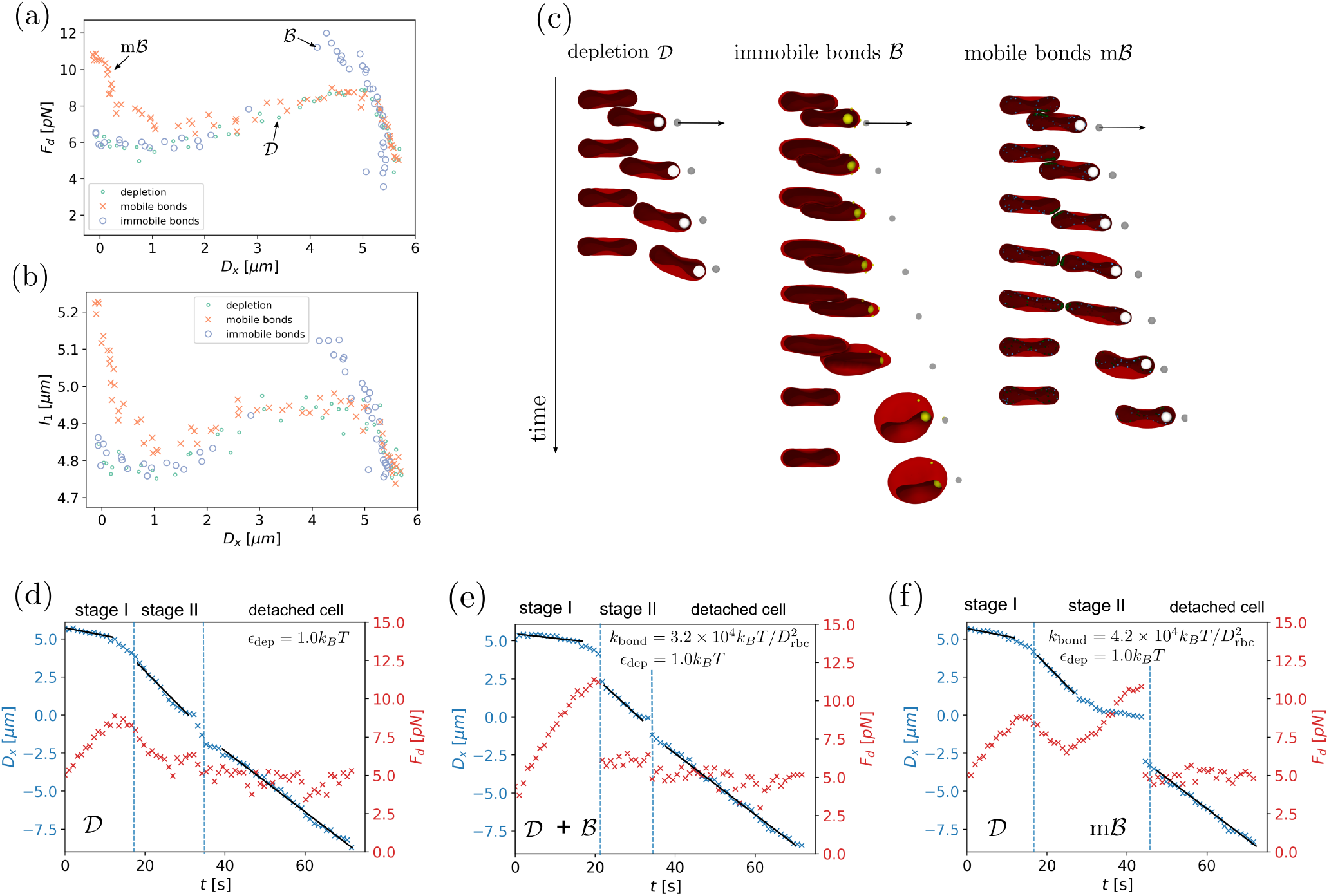
Comparison of RBC disaggregation for different adhesion models (*ϵ*_dep_ = 1*k*_*B*_*T* in all cases). (a) Disaggregation force *F*_*d*_ and (b) RBC extension characterized by *I*_1_ as a function of *D*_*x*_. RBCs are initially placed with a 60% overlap. (c) Snapshots of the RBCs for different times and aggregation models, see also Movies S2-S4 [41]. Disaggregation process as a function of time for the three models: (d) the *D* model, (e) the *B* model, and (f) the *mB* model. Two disaggregation stages can be defined, including initial yielding with nearly no change in *D*_*x*_ (stage I) and motion of the RBCs relative to each other (stage II).

In conclusion, our comparison of simulations and experiments of RBC disaggregation strongly suggests the presence of both depletion and bridging interactions in RBC aggregation. The initial increase in *F*_*d*_ is due to depletion interactions, while the second increase in *F*_*d*_ is mediated by mobile bridges. In fact, when the time of initial contact of two RBCs in experiments is short, there is nearly no increase in *F*_*d*_ near the time point of final RBC separation. Therefore, the formation of bridges requires some contact time for two cells. Our simulations also demonstrate that formed bridges must be mobile, since immobile bonds generally lead to an abrupt separation of two RBCs before they even reach the edge-to-edge configuration. This would be consistent with weak adsorption interactions of bridging molecules with the glycocalyx layer at the RBC surface, which allows sliding motion. Despite a favorable comparison between experiments and simulations and the fact that the presented models implement the suggested mechanisms (depletion and bridging) for RBC aggregation, we cannot fully exclude a possibility for the existence of another aggregation model. Furthermore, a better characterization of depletion and bridging contributions to RBC aggregation requires careful experiments, where depletion and bridging interactions can be controlled independently. For example, depletion interactions without bridging contribution were mediated through the concentration of rod-like *fd*-viruses in a recent study [30]. Such a control of aggregation interactions between RBCs would allow a quantitative comparison between simulations and experimental measurements.

## Supporting information

Experimental and simulation comparison

simulation video for depletion only interaction

simulation video for immobile bridges interaction

simulation video for mobile bridges interaction

## ACKNOWLEDGMENTS

We thank Kisung Lee (Institute of Basic Science, South Korea) for stimulating discussions. D.A.F acknowledges computing time on the supercomputer JURECA [52] at Forschungszentrum Jülich under grant no. actsys. N.M and M.E. acknowledge the support of the Basque Government through the BERC 2022-2025 program and the Ministry of Science and Innovation: BCAM Severo Ochoa accreditation CEX2021-001142-S / MICIN / AEI / 10.13039/501100011033. C.W acknowledges the support from the German Research Foundation (DFG FOR 2688, Projects No. WA 1336/12-2. N.M. acknowledges the support from the European Union’s Horizon 2020 under the Marie Sklodowska-Curie Individual Fellowships grant 101021893, with acronym ViBRheo.

## AUTHOR CONTRIBUTIONS

N.M. and D.A.F. designed the research project. N.M. performed the simulations and analysed the obtained data. A.T. helped in creating the simulation setup. K.K., A.S., and T.J. performed experiments and analysed the data. All authors participated in the discussions and writing of the manuscript.

