## Supplementary Information for "Aggregation and disaggregation of red blood cells: depletion versus bridging"

### I. LASER TWEEZERS CALIBRATION

The main steps of laser tweezers calibration described in Ref. [1] are as follows. In the presence of external force and the associated potential $U(x)$, the equilibrium probability density distribution is given by

$$\text{pdf} \propto e^{-\frac{U(x)}{k_B \cdot T}}, \tag{1}$$

where $k_B$ is the Boltzmann constant and $T$ is temperature. The potential $U(x)$ from the optical trap is harmonic

$$U(x) = \frac{1}{2} k_x (x - x_0)^2, \tag{2}$$

where $k_x$ is the stiffness of optical trap in $x$ direction and $x_0$ is the equilibrium position of the bead in the trap.

According to the equipartition theorem, the energy, which corresponds to each coordinate, is on average

$$\langle U(x) \rangle = \frac{k_B T}{2}, \tag{3}$$

and the functional dependence between the probability and the bead position in the optical trap can be described by a Gaussian distribution as

$$f(x) \propto e^{-\frac{1}{2}\left(\frac{x-x_0}{\sigma_x}\right)^2}, \tag{4}$$

where $\sigma_x$ is the standard deviation. Considering Eqs. (1-4), we can write the following

$$k_x = \frac{k_B T}{\sigma_x^2}. \tag{5}$$

To estimate $\sigma_x$, a single microbead is trapped by optical tweezers, and its movement inside the trap is recorded with a frame rate of 100 frames per second. Figure 1 shows probability density functions (one- and two-dimensional) of the position of trapped bead, from which $\sigma_x$ is computed. Then, $k_x$ is given by Eq. (5) for a fixed laser power. This experiment is performed for several different laser powers, leading to the calibration curve shown in Fig. 2, from which the stiffness of the trap is fitted as

$$k_{trap} \left[\frac{pN}{\mu m}\right] = k_x \left[\frac{pN}{\mu m}\right] = a \cdot P\,[mW]\,, \tag{6}$$

where $a = (576 \pm 16) \cdot 10^{-4}\, pN/(\mu m \cdot mW)$.

*

†

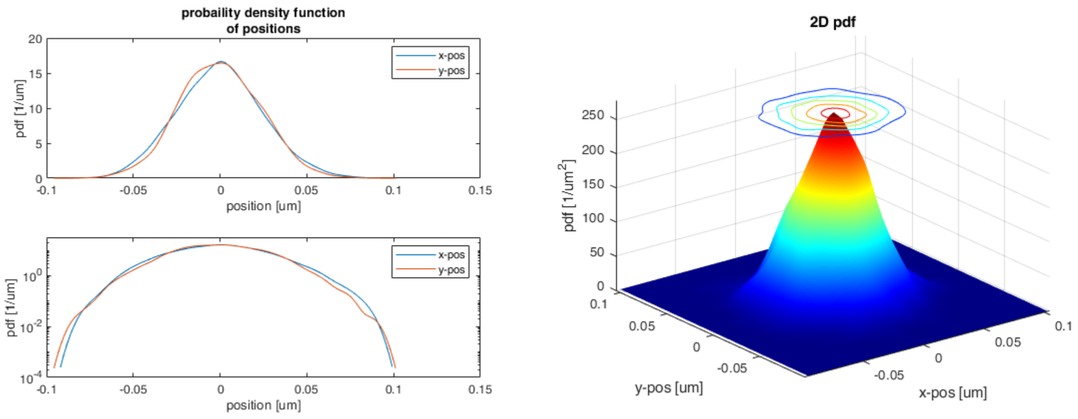


FIG. 1. Probability density function of a bead position (a) for a single coordinate and (b) in two-dimensional representation.

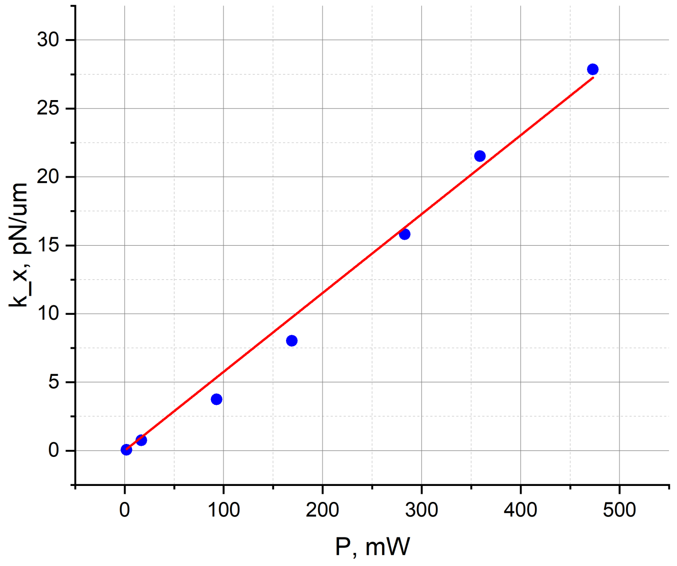


FIG. 2. Optical tweezers calibration curve, showing the dependence of the trap stiffness on the laser power.

### II. EFFECT OF INITIAL CONTACT TIME ON THE NUMBER OF BONDS

For the $m\mathcal{B}$ model, the number of formed bonds is affected by the time of initial contact and the surface density $\rho_{\rm sites} = n_b/A_{\rm rbc}$ of binders, because the binders diffuse inside the RBCs and can form bonds only when they are close enough to a free binding partner. Figure 3

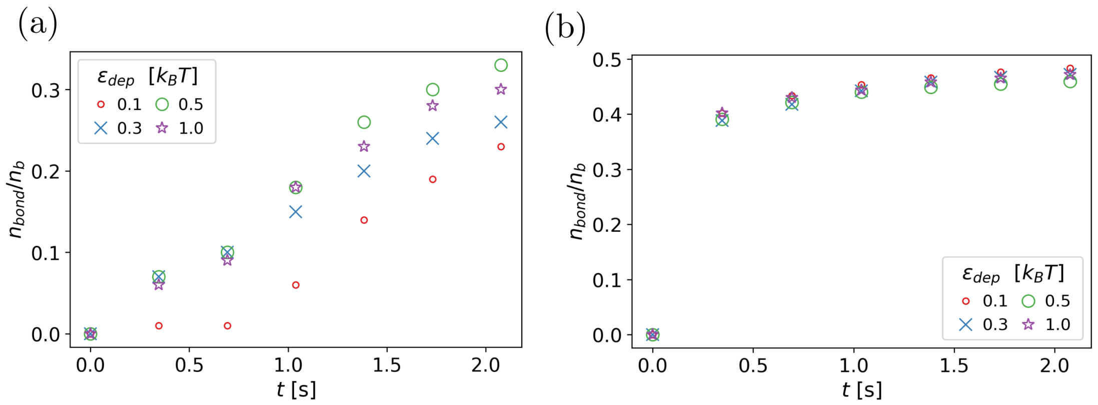


FIG. 3. Initial aggregation of two RBCs with an overlap of 60% for the $m\mathcal{B}$ model. Number $n_{bond}$ of formed bonds normalized by the number $n_b$ of binders as a function of time for (a) $\rho_{\rm sites} = 106/A_{\rm rbc}$ and (b) $\rho_{\rm sites} = 1463/A_{\rm rbc}$, and various depletion strengths $\epsilon_{\rm dep}$.

shows the number $n_{bond}$ of formed bonds as a function of time for two different densities of binders. We observe that $n_{bond}$ saturates after a contact time of about 2 s for various depletion strengths $\epsilon_{\rm dep}$. Depletion interactions aid the formation of bonds for the low density of binders, since they force the RBCs to stay in a close contact. For the large $\rho_{\rm sites}$ in Fig. 3(b), the effect of depletion on $n_{bond}$ is negligible, because a few bonds form quickly, keeping the two RBCs in a close contact. As a result, we let the RBCs to initially aggregate for 2 s, to avoid the influence of initial contact time on the measurements of disaggregation force.

### III. DESCRIPTION OF MOVIES

**Movie S1:** Side-by-side simulation and optical tweezers experiment of the disaggregation of two RBCs. The simulation is conducted using the $m\mathcal{B}$ model with $\epsilon_{\rm dep} = 1.3k_BT$ and $k_{\rm bond} = 3.3 \times 10^4 k_BT/D_{\rm rbc}^2$. In the simulation, a cross-section of the RBCs is shown. Green particles represent the formed bridges, whereas white and gray particles mimic the pulling force exerted by optical tweezers. The experimental setup corresponds to a RBC doublet in plasma, using a bead-assisted disaggregation scheme.

**Movie S2:** Disaggregation of two RBCs for the $\mathcal{D}$ model, see Fig. 7(c) in the main text. The simulation is conducted using the $\mathcal{D}$ model with $\epsilon_{\rm dep} = 1.0k_BT$. A cross-section of the RBCs is shown, where white and gray particles mimic the pulling force exerted by optical tweezers. The white particle is linked to the gray particle via a harmonic potential, and the gray particle is pulled at a constant rate.

**Movie S3:** Separation of aggregated RBCs for the $\mathcal{B}$ model, see Fig. 7(c) in the main text. The simulations is performed using the $\mathcal{B}$ model with $\epsilon_{\rm dep} = 1.0k_BT$ and $k_{\rm bond} = 3.2 \times 10^4 k_BT/D_{\rm rbc}^2$. A cross-section of the RBCs is shown, where white and gray particles mimic the pulling force exerted by optical tweezers.

**Movie S4:** Disaggregation of two RBCs for the $m\mathcal{B}$ model, see Fig. 7(c) in the main text. The simulations is conducted using the $m\mathcal{B}$ model with $\rho_{\rm sites} = 106/A_{\rm rbc}$, $\epsilon_{\rm dep} = 1.0k_BT$, and $k_{\rm bond} = 4.2 \times 10^4 k_BT/D_{\rm rbc}^2$. A cross-section of the RBCs is shown, where green particles represent the formed bridges, light blue particle are free binders (not bound), and white and gray particles mimic the pulling force exerted by optical tweezers.

[1] Pesce, Giuseppe, Jones, Philip H., Maragò, Onofrio M., and Volpe, Giovanni, Optical tweezers: theory and practice, Eur. Phys. J. Plus **135**, 949 (2020).
